# Microbial redox activity mediated anaerobic pyrite oxidation under circumneutral conditions

**DOI:** 10.1101/2020.02.17.952168

**Authors:** Tong Liu, Yutian Hu, Nan Chen, Linlin Ma, Qiaochong He, Chuanping Feng

## Abstract

In modern Earth, anaerobic pyrite oxidation under circumneutral conditions also has great impact on the fate of nitrate in aquifers and sediments, as well as the transportation of toxic metals. However, the mechanism of how microbes mediated this process is still being debated. Electrochemical analysis on pyrite cubic electrode showed that, its oxidation threshold under anaerobic circumneutral conditions (ca. 200 mV) was much lower than that at aerobic acidic conditions (ca. 650 mV), implying possible direct pyrite oxidation by high redox potential cellular components. Sole substrate (pyrite) microbial enrichment cultures with EDTA addition showed higher oxidation rate (0.092 d^-1^) than that of EDTA-free cultures (0.019 d^-1^), suggesting that ligands producing pathway was much preferred by microbes than maintaining acidic micro-environments. This hypothesis was supported by amplicon and metagenomic sequencing data, which demonstrated discrepant bacteria involving iron-sulfur oxidation and metabolic potentials in cultures with/without EDTA addition. A concept model was proposed based on experimental data considering different reaction stages and microbial communities. The results shed lights on the potential interactions between microbes and pyrite, which may serve as a model for explaining subsurface pyrite oxidation and optimizing anaerobic pyrite oxidation-based pollutant removal processes.

**TOC art:** 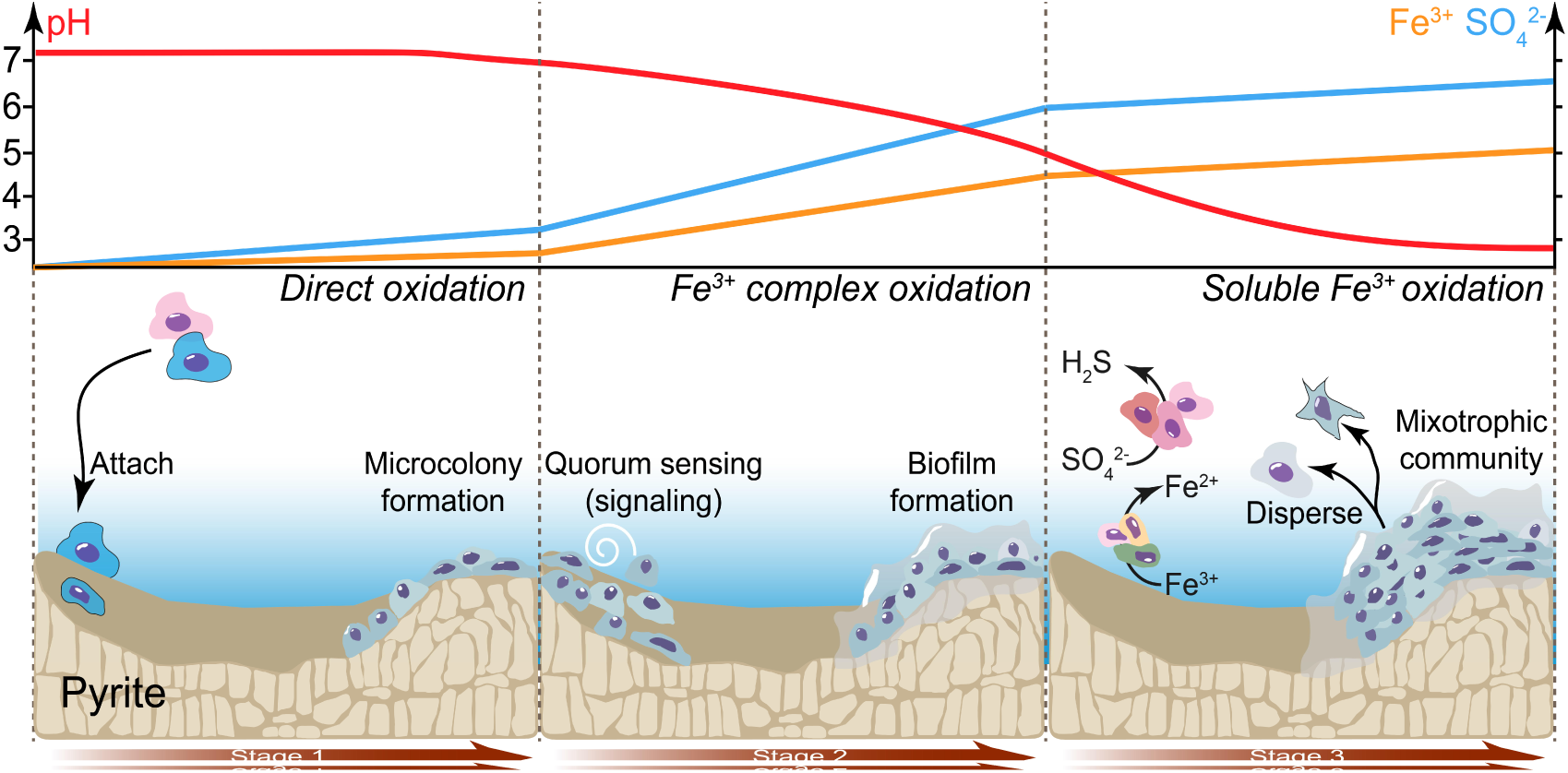

## Introduction

Pyrite (FeS_2_) is by mass the most abundant thermodynamically stable iron-sulfide mineral in earth’s crust ^1^. Over geological times, its burial and weathering interreacted intensively with organic matter in sediments and oxygen in atmosphere ^2^, thus played a key role in the variation of earth’s surface chemical system (e.g. the Canfield ocean ^3^) and the evolution of life diversity ^4^. For modern earth, despite the importance of pyrite oxidation on earth’s element cycling, the transportation of contaminants and formation of acid mine drainage (AMD) are also strongly affected ^5^. Microbial pyrite oxidation has been studied extensively under aerobic conditions with low pH, typically in the context of AMD formation. Acidophilic microbes, represented by *Acidithiobacillus ferrooxidans*, prominently accelerates pyrite dissolution through aerobic soluble Fe^2+^ oxidation, which produces soluble Fe^3+^ as chemical oxidant for pyrite ^6,7^. However, for most geological sites, such as soil, sediments, and shallow aquifers, anaerobic and buffered geochemical conditions, are more common than an aerobic environment with low pH ^8^.

The simultaneous nitrate reduction and sulfate production has been observed at multiple pyrite-bearing aquifers over the past few decades ^9,10^. A series of field study suggested that the coupling of microbial mediated chemolithotrophic denitrification and pyrite oxidation contributed to the nitrate consumption, sulfate production ^9,11,12^, release of trace elements in pyrite lattice ^13,14^, and stable isotope fractionation of ^15^N, ^18^O, and ^34^S ^15,16^. Laboratory experiment using pyrite nanoparticles and *Thiobacillus denitrificans* (*T*.*denitrificans*) provided direct evidence of simultaneous nitrate reduction and ferric and sulfate production ^17^, where an electron balance considering the formation of Fe^3+^ and sulfate along with the transformation of nitrate to dinitrogen was in consistence with the stoichiometry (Eq. 1).

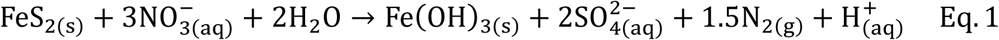

Nevertheless, as an acid-non-soluble metal sulfides, pyrite derived all valance bands from metal orbitals, leading its crystal substantial against proton promoted or direct microbial dissolution ^18^. Consequently, the role of functional microbes and the mechanism of the chemical variations in pyrite-bearing aquifer are not clear ^19^. In addition, anaerobic pyrite oxidation process was recognized as promising method in remediating nitrate contamination ^20–23^. However, the circumneutral condition limited pyrite oxidation process, resulting in low nitrate removal efficiency ^24–26^. which in turn called for fundamental mechanism studies.

On the other hand, research of pyrite oxidation under AMD-like conditions has dived far into molecular and genetic level ^27^. The utilization of state-of-art techniques, such as corrosion electrochemistry and molecular biology, enables the understanding of the mechanism and kinetics of this process from the mineralogy aspects ^18,28,29^ and the means of process acceleration by microbes ^30^.. Two possible pathways drive the anaerobic pyrite oxidation at neutral pH ^31^. In the indirect pathway, pyrite is attacked by chelated Fe^3+^, which requires either an acidic nano-environment or putative chelated compounds, microbes therefore acquire energy through iron assisted electron-shuttling.

Alternatively, pyrite may be oxidized via direct microbial attack along with the cell colonization, which is postulated need the participation of putative cell compounds. Although both pathways may explain the role of microbes, more experimental evidence are required to validate the pathways exist or not and to determine which pathway excavate more electrons from pyrite oxidation for microbes.

The goal of this research is to elucidate the mechanism of microbial mediated anaerobic pyrite oxidation under neutral pH and to promote optimization of pyrite oxidation-based pollution control in municipal plants or aquifers. We analyzed electrochemical oxidation behavior using pyrite cubic electrode and conducted microbial experiments with pyrite as sole substrate and nitrate as electron accepter to elucidate if microorganisms can oxidize pyrite directly and which pathway dominated pyrite oxidation. The results obtained not only enhances our understanding of the role microbe plays while interreacting with solid pyrite, but also has potentially broad environmental impacts and implications for major elements cycling from both geochemistry and environmental remediation aspects.

## Results

### 1. Redox potential limiting anaerobic microbial pyrite oxidation

Of the electrochemical mixed potential theory (Figure 1a), the mixed potential (E_m_) referred to the measured redox potential difference between anode and cathode, which was the energy pushed the electrochemical reaction. E_m_ was also equal to open circuit potential (E_ocp_) that no redox reaction happens between electrodes and redox active species in solution. E_ocp_ analysis of pyrite electrode showed an intense decline from 0.42 to 0.10 V (vs. standard hydrogen electrode (SHE)) with the increase of electrolyte pH from 1 to 11, and E_ocp_ altered little within the pH range of 5-9 (Figure 1b). Similarly, linear sweep voltammetry evaluation showed decreased pyrite electrode stability with electrolyte accretion (Figure 1c), the asymmetric curves of Tafel plots (Figure 1c) also suggested that the pyrite electrode was oxidized irreversibly.

**Figure 1.**
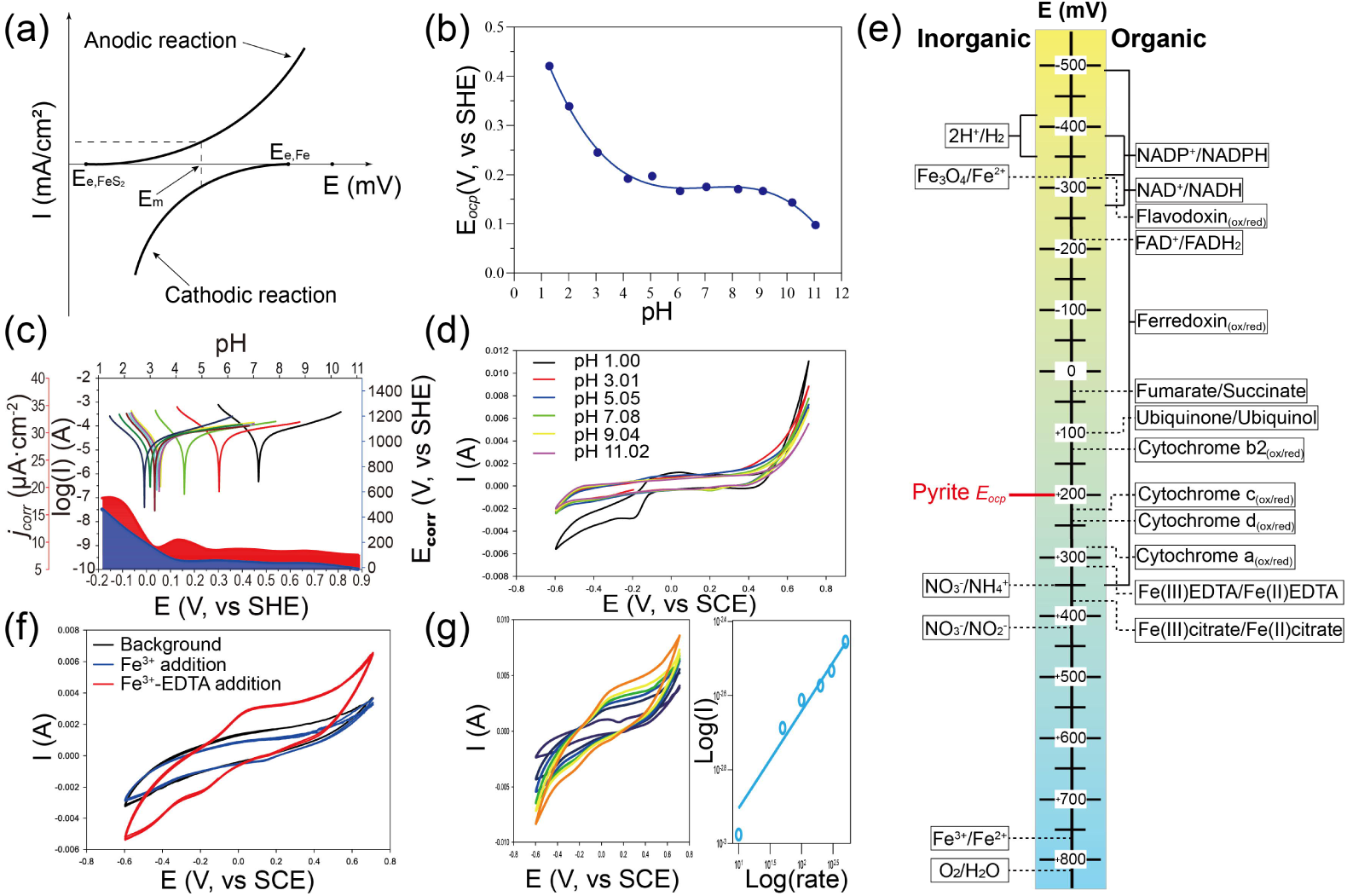
Electrochemical profile of anaerobic pyrite oxidation under neutral conditions using lab-made pyrite cubic electrode. Schematic illustration of electrochemical mixed potential theory (a), E_ocp_ variation in 0.1 M Na_2_SO_4_ solution with various pH (b), Tafel plot for pyrite electrode in 0.1 M Na_2_SO_4_ solution with various pH (c), Cyclic voltammetry curves of pyrite electrode in 0.1 M Na_2_SO_4_ solution with different pH (d), Redox potentials (vs. SHE) of important redox reactions in electron transport chains in cell (Modified from ^32^) (e), Cyclic voltammetry curves of pyrite electrode in 0.1 M Na_2_SO_4_ solution (pH 7.0) with Fe^3+^ or Fe^3+^-EDTA addition (f), Successive cyclic voltammetry scans of pyrite electrode in 0.1 M Na_2_SO_4_ solution (pH 7.0) with Fe^3+^-EDTA addition with increased scan speeds and log/log plot of change in anodic peak currents vs. change in scan rate (g).

When cyclic voltammetry (CV) was performed in anaerobic electrolyte, the change of electrolyte pH did not alter the reactivity of pyrite electrode (Figure 1d). Because soluble Fe^3+^ only presented in pH 1 condition, its reduction to Fe^2+^ on pyrite surface was verified with the significant catholic peak observed (Figure 1d). Based on the E_ocp_ data, which was confirmed by linear sweep and cyclic voltammetry, a redox potential ladder under physiological pH (Figure 1e) was constructed from previous data ^32^. Fe^3+^-EDTA was able to oxidize pyrite under neutral pH (Figure 1e), which was tested by CV scan with Fe^3+^ or Fe^3+^-EDTA added in pH 7 electrolyte. We observed significant anodic and catholic peaks (Figure 1f) when adding Fe^3+^-EDTA, indicting Fe^3+^-EDTA can oxidize pyrite under neutral pH, while Fe^3+^ not. To distinguish whether diffusion or adsorption was the rate-limiting step of redox active molecules’ (i.e. Fe^3+^-EDTA) reactions on electrode surface, we increased the CV scan rate gradually from 10 to 2,000 mV/s (Figure 1g), and an increase in peak current was observed. Molecules may be oxidized/reduced on the electrode surface by diffusion or adsorption controlled mechanisms ^33^, which can be distinguished by successive CV scan with gradually increased scan rate ^34^. Given current rise was proportional to the square root of the scan speed in diffusion-controlled process ^35^, a diffusion-controlled mechanism was confirmed in the case of Fe^3+^-EDTA on pyrite surfaces, in which a slope of 0.26 with *R*^*2*^ = 0.971 was obtained by plotting a log(I) (where I is the anodic peak current) against log(rate) (where rate is the scan rate in mV/s) (Figure 1g).

### 2. Accelerated anaerobic microbial pyrite oxidation by EDTA addition

Without any lag period, nitrate concentration in experiments with pyrite and microorganisms decreased rapidly in the first 3 days after inoculation, and the decrease rate slowed down thereafter (Figure 2a). However, significantly faster nitrate removal rate was observed in experiments with EDTA addition (Rec, *p* = 0.0078), nearly complete nitrate removal was achieved within 25 days, while only half nitrate was removed without EDTA addition (Ctrl). Similarly, nitrate reduction rates for treatments without inoculum or growth substrate (i.e. pyrite) was much smaller than EDTA amended microcosm set. Correspondingly, nitrite accumulation rate was much higher than the Ctrl. The peak nitrite concentrations of Rec reached 18 mg-N/L on day 4, while the peak nitrite concentration of Ctrl did not show up until day 15 (∼6 mg-N/L).. Ammonia was not detected in all treatments. Total iron concentration in Rec had a similar trend as nitrate, of which the concentration increased rapidly to ca. 30 mg/L in the first 5 days, and kept a slow increment thereafter (Figure 2c). Obvious augment of total iron was observed only in Rec (Figure 2c), while for other treatments, the total iron concentrations were below 2 mg/L.. In addition, ferrous iron concentrations in all treatments were always lower than 1 mg/L in all experiments (Figure 2d), indicating ferric iron was the major iron specie.

**Figure 2.**
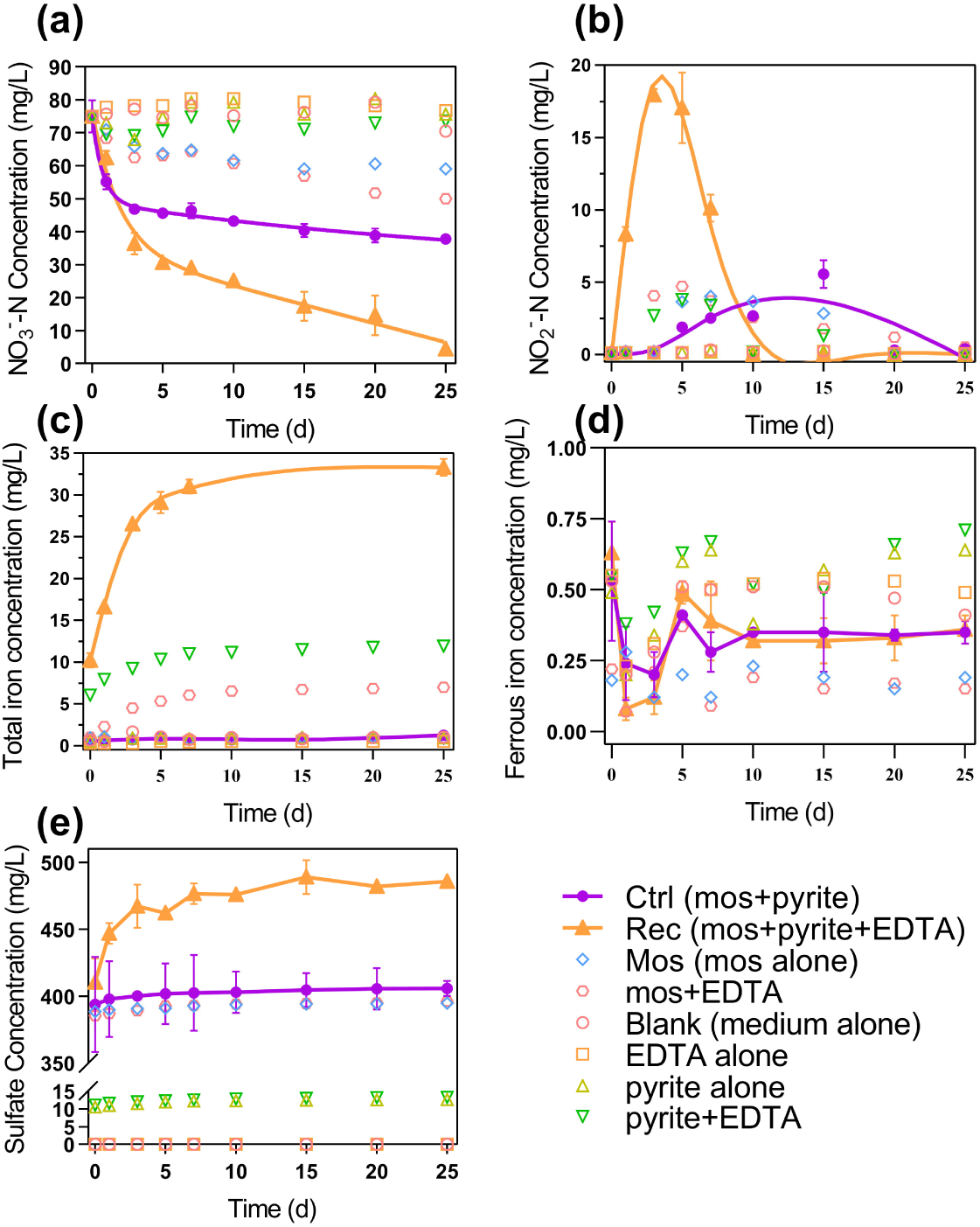
Changes in aqueous species concentrations during microbial mediated anaerobic pyrite oxidation under circumneutral pH. NO_3_^-^ (a), NO_2_^-^ (b), total iron (c), ferrous iron (d) and SO_4_^2-^ (e) concentration variation under experimental conditions with/without EDTA-addition and microorganisms (mos) inoculation.

Despite trace amount of reduced sulfur compounds from inoculum, pyrite was regarded as the major electron donor for microbes. After 25 days of experiment, the increment of sulfate concentration was 12.04, 73.96, 6.91, and 2.13 mg/L for Ctrl, Rec, Mos, and abiotic control, respectively (Figure 2e). Only a little sulfate was produced in abiotic control, therefor it was confirmed the abiotic pyrite oxidation process can be neglected. Theoretical mass ratio (w/w) of the produced sulfate and consumed nitrate was calculated to be 1.03 based on Eq. 1. The actual mass ratio for Rec and Ctrl was 1.07 and 0.32, respectively. While a small mass ratio deviation was found for Rec (3.9%), mass ratio for Ctrl varied a lot (68.9%). The mass ratio deviation for Ctrl was ascribed to the cooccurrence of autotrophic process using pyrite as electron donor and heterotrophic process with necromass as electron source under low-energy-supply conditions (Figure 2a), since the heterotrophic process could compete with autotrophic process for electron accepter (i.e. nitrate). Compared with high sulfate background concentration, a low sulfate concentration produced in Ctrl may also hinder the mass ratio calculation.

### 3. Microbes tend to gather onto pyrite surface

The living state (i.e. planktonic or biofilm) of microbes has a strong impact on their interactive relationship with pyrite, thus the supernatant of microbial culture and the cell-mineral association were assayed. 3D-EEM (three-dimension excitation emission matrix) of the supernatant from cell + pyrite cultures showed the lowest fluorescence intensity (Figure 3a, Figure S4), implying potential microbial accumulation near pyrite particles. Meanwhile, EDTA addition suppressed biofilm formation ^36^, leading to a multiplied cell number in cultures with EDTA (Figure 3a, Figure S3). The microbial accumulation near pyrite and on pyrite surface were further verified using fluorescence microscopy, CLSM (confocal laser scanning microscope), and ESEM (environmental scanning electron microscope) (Figure 3b-d). In cultures without EDTA, microbes associated with pyrite particles and formed compacted cell-mineral complex. While in cultures with EDTA addition, the number of cells on pyrite surface decreased, which agreed with the 3D-EEM data of supernatant.

**Figure 3.**
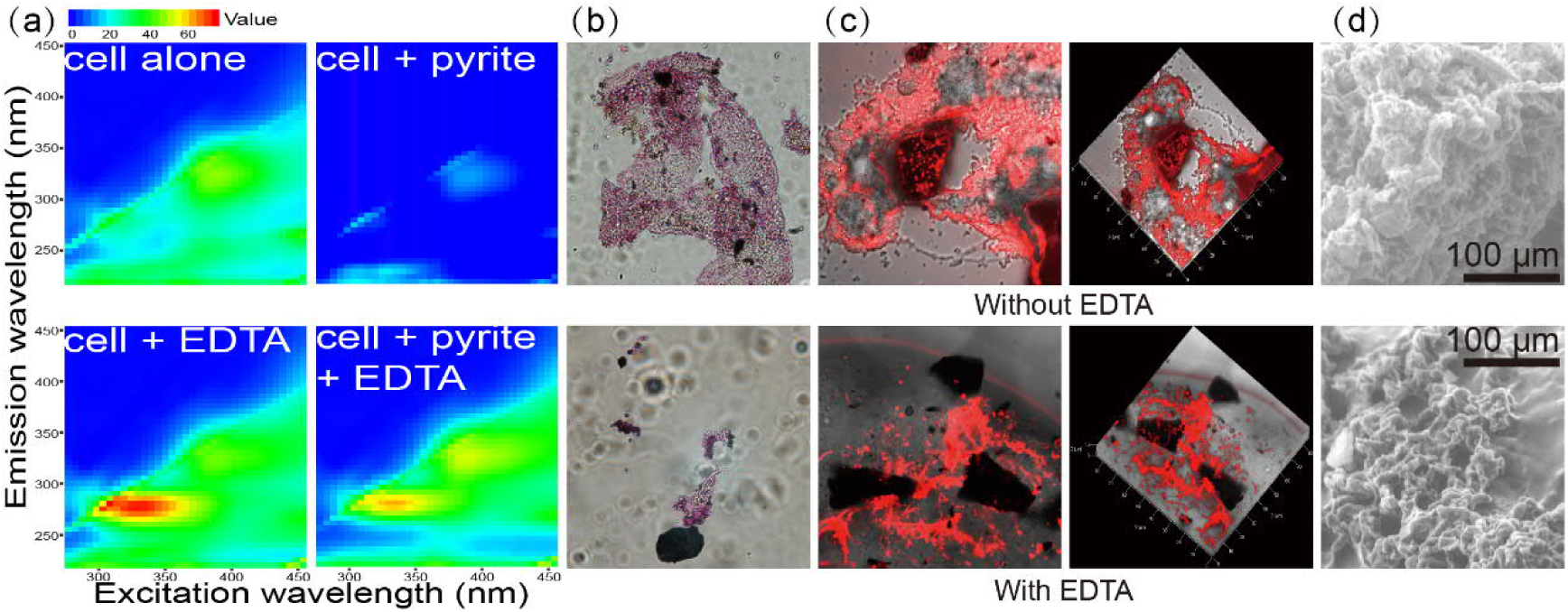
Microbial cultures tend to associate with pyrite surfaces. 3D-EEM spectra of the medium supernatant after ca. 10 days of growth (a). DiI cell membrane stain (red) and fluorescence images of the microbial cultures (b). DiI cell membrane stain and CLSM images of the cell-mineral association from parallel microbial cultures after ca. 10 days of growth (c). ESEM images of the cell-mineral association at same magnification (d).

### 4. Microbial community analysis

The taxonomic composition of microbial cultures based on amplicon sequencing showed that microbial communities were dominated by phylum Bacteroidetes and Proteobacteria with lower abundance in microbial cultures without EDTA (Figure 4a). However, phylum Proteobacteria got more dominative in cultures with EDTA addition (Figure 4a). At genus level (Figure 4b, Figure S12), sulfur and nitrogen cycles related genera (*Sulfurimonas, Denitrobacter*, and an unassigned genus from family *Desulfobacteraceae*) showed a relatively higher abundance in EDTA-addition culture, while in EDTA-free culture the high abundance genera switched to *Marinobactor* and *Thiobacillus* (Figure 4b). Furthermore, genera B-42, *Paludbacter*, and *Bacillus* tend to govern the starved culture (Mos).

**Figure 4.**
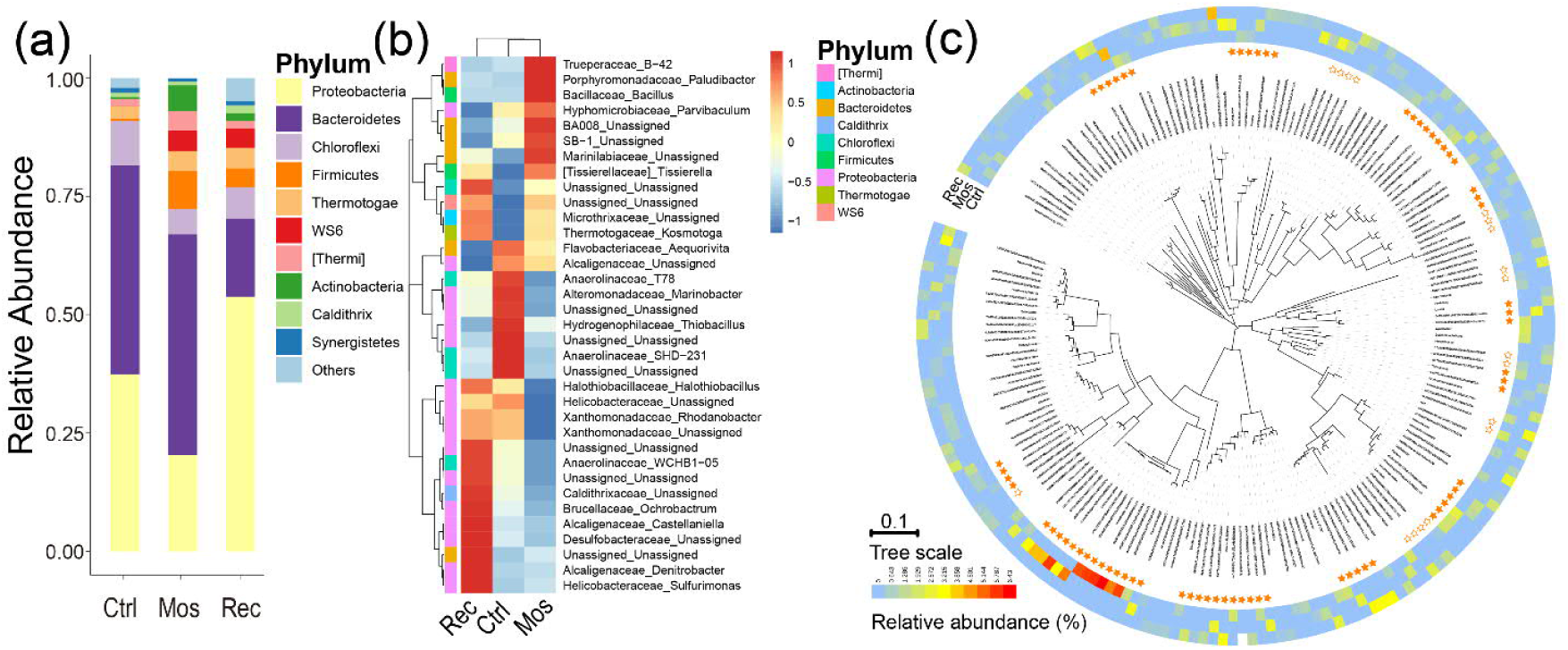
Microbial community structure based on 16S rRNA gene amplicon sequences. The stacked bar plot shows the relative abundances of top 10 phyla (a). Relative abundance of the top 35 genera (b). Phylogeny, abundance, and core microbiome of bacterial ASVs in the 3 microbiotas (c) (interactive phylogenic tree can be found at https://itol.embl.de/tree/602475147460471565937282).

Taxonomic binning of the 3 metagenomes yielded 87 high quality (genomic completeness higher than 90% and contamination lower than 5%) metagenomic assembled genomes (MAG) (Figure 5a). MAGs recovered from Rec had larger genome size (Figure 5a), which indicated a higher evolution complexity and functional redundancy ^37^. The MAGs obtained were then annotated with eggNOG database ^38^ and the genes were mapped to KEGG database ^39^. The abundance of genes involved in major energy metabolism pathways showed that MAGs from Rec had the highest abundance of energy metabolism genes in every sub-pathway (Figure 5b). Genes related to cellular community and mobility demonstrated that more genes in Ctrl MAGs involved in bacterial chemotaxis and flagellar assembly process (Figure 5c), which suggested potential cell movement towards solid substrate (i.e. pyrite). While Rec MAGs had the highest abundance of quorum sensing encoding genes (Figure 5c), which suggested a more complexed microbial community.

**Figure 5.**
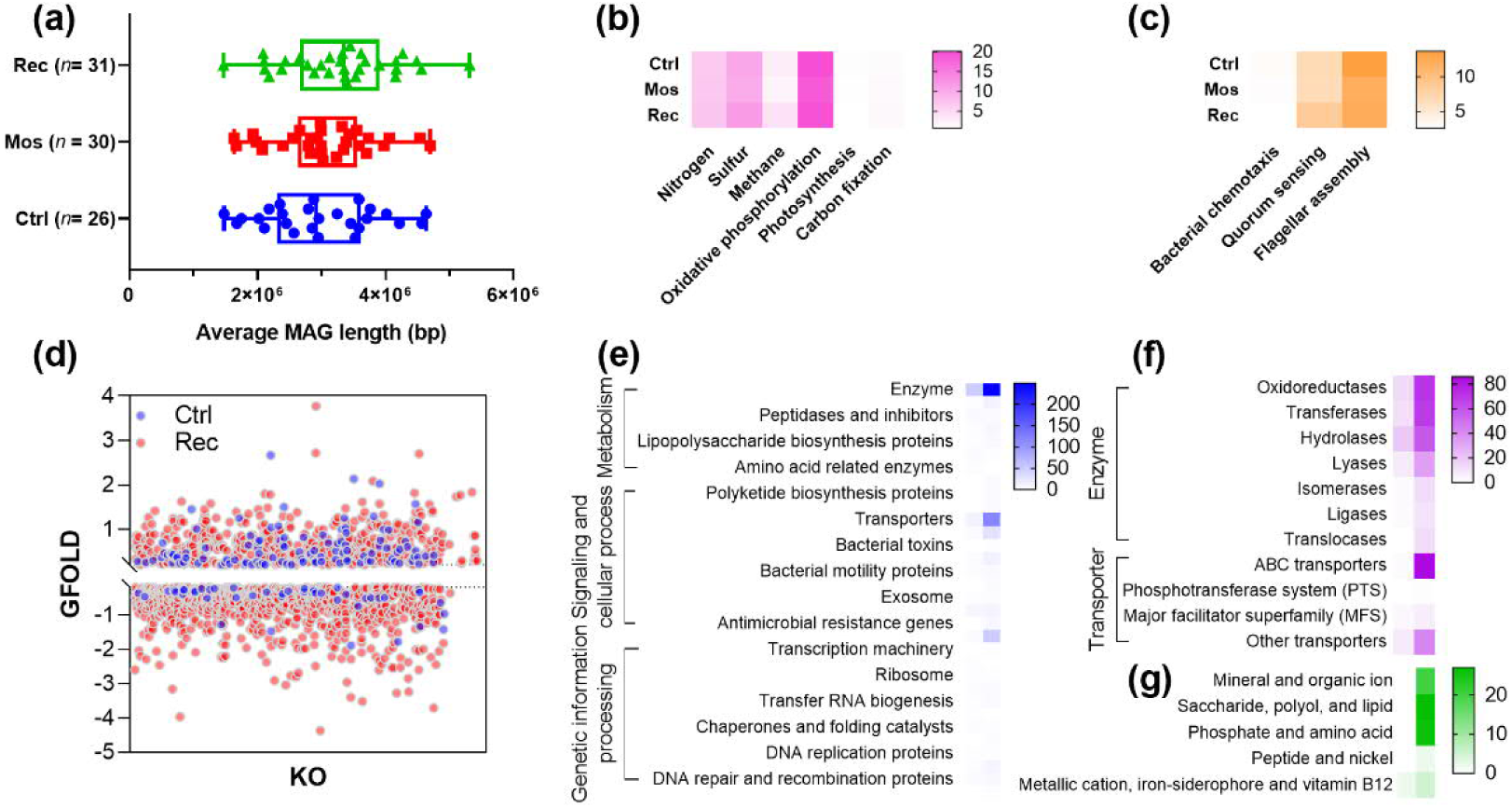
Functional structure based on metagenomic sequencing. MAGs from cultures without EDTA addition had smaller MAG length (a). EDTA-addition cultures had higher abundance of energy metabolism genes, legend (genes/genome) (b). Cellular community and mobility related genes increased with more energy available, legend (genes/genome) (c). Compared with EDTA-free cultures, EDTA addition cultures had more discrepant genes, indicating an oligotrophic state of EDTA-free cultures (d). Based on KEGG protein family analysis, high abundance genes of Rec cultures mainly involved those encoding enzymes and transporters (e). High abundance enzyme encoding genes of Ctrl and Rec culture were of different kinds and abundance, and Rec culture had the most abundant of ABC transporters encoding genes (f). Within ABC transporters encoding genes, organic compounds transportation related genes were of high abundance in Rec culture, while high abundance genes in Ctrl cultures involved mainly in metallic, cation, iron-siderophore, and vitamin B12 transportation (g).

To evaluate the functional difference between cultures Rec and Ctrl, the differential abundance of the annotated genes from these two metagenomes were calculated using Mos metagenome as reference (Figure 5d). Under a differential threshold of 0.2 GFOLD, more genes with higher abundance were found from Rec metagenome (Figure 5d), implying an immense functional diversity. While the minor difference between Ctrl and Mos metagenomes (Figure 5d) indicated their similarity in metabolic potential. The high abundance differential genes were further analyzed with KEGG protein family (Figure 5e), which showed, in both Ctrl and Rec metagenomes, genes encoding enzymes and transporters were of higher abundance, and the gene abundance of Rec are higher than Ctrl. Among the enzyme encoding genes, only hydrolases were of high abundance in Ctrl, while Rec had several high abundance enzymes (Figure 5f). Meanwhile, ABC transporters encoding genes of Rec were of the highest abundance (Figure 5f). It was noteworthy that several organic compounds transportation (such as saccharide, lipid, and amino acids) related ABC transporters were of high abundance in Rec, while only genes involved in metallic, cation, iron-siderophore, and vitamin B12 transportation were of high abundance in Ctrl (Figure 5g). From the numbers of differential genes (Figure 5d) and the similar abundance pattern across different levels of protein families (Figure 5e-g), genes from Rec were always of higher abundance in more functional categories. Thus it was concluded that the microbial community in Rec culture was of higher functional diversity and redundancy.

## Discussions

### 1. Microbes could oxidize pyrite by either cellular compounds directly or Fe^3+^-complex

The E_ocp_ of pyrite electrode was examined to decrease with the pH increment under anaerobic conditions (Figure 1b), which was in analogous to that observed in aerobic basic ^40^ and acidic situations ^41^. Since the rate-limiting step of aerobic pyrite oxidation was the electron transfer from pyrite bulk crystal to surface adsorbed O_2_ molecular, the decreased E_ocp_ was attributed to reduced obstacle of electron transfer ^42^. The measured E_ocp_ was ca. 0.2 V at pH 7.0 (Figure 1b), which was higher than that from literature (ca. −0.28 V) ^43^. However, the E_ocp_ that Tao et al. ^43^ obtained varied a lot among different samples, and discrepant electrode preparation methods and electrolyte may also arise this issue. Even though the measured E_ocp_ value varied, its tendency was reliable, and was also validated by the corrosion current decline in Tafel assay (Figure 1c). Under aerobic acidic situations, pyrite E_ocp_ was reported to be ca. 0.6 V ^44,45^, and pyrite oxidation can merely happen under redox potential of 0.65 V ^46^. It can therefore be concluded that E_ocp_ was a key discrepancy between acidic and basic cases. Furthermore, CV analysis did not show obvious pyrite oxidation at pH above 2.0 (Figure 1d), while unambiguous oxidation occurred at pH 2.0. Pyrite oxidation at pH 2.0 was in consistence with previous study ^45^, and can be interpreted with the enhanced solubility of Fe^3+^.

To demonstrate the relationship between pyrite E_ocp_ and microbial mediated redox reactions under circumneutral conditions, the pyrite E_ocp_, redox potentials of common chemical reaction, and potentials of important cellular redox reactions were summarized into a ladder plot (Figure 1e). Many cellular compounds were then able to oxidize pyrite directly thermodynamically, which was of immense disparity with the fact that pyrite cannot be oxidized directly by cellular compounds under acidic conditions ^46^. Previous study showed there were reversible redox reactions between pyrite and hydroquinone molecules ^33^, which was in agreement with direct pyrite oxidation (Figure 1e). Considering the rate-limiting step in hydroquinone and pyrite interaction was hydroquinone diffusion rather than interfacial electron transfer ^33^, the direct pyrite oxidation by cell components was further supported. For the direct and indirect pathways of anaerobic pyrite oxidation under circumneutral pH ^31^, it was not reasonable for microbes maintaining an acidic microenvironment. Because the pyrite E_ocp_ increased greatly with pH dropdown (Figure 1b), obtaining high redox potential compounds with acidic environments was not energy efficient for microbes. On the other hand, high redox potentials can be easily obtained with ligands like EDTA or citrate (Figure 1e), thus making the indirect pathway much convincing. To validate indirect pyrite oxidation, CV assay was carried out with EDTA addition, and we observed significant pyrite oxidation (Figure 1f). The Fe^3+^-EDTA complex diffusion limited the overall reaction indicated a fast surface electron transfer (Figure 1g), which may further support the hypothesis that microbes accelerated pyrite oxidation by producing ligands.

Respecting microbes may produce ligands to accelerate pyrite oxidation, it was reasonable to hypothesis that pyrite oxidation can be strengthened by adding ligands. By adding EDTA, pyrite oxidation was enhanced significantly from 0.019 to 0.092 d^-1^ (Figure 2). The pyrite dosage normalized pseudo-first order kinetics (0.009 L/(g·d)) was also near 2 times greater as that (0.005 L/(g·d)) of previous study in our laboratory using similar inoculum ^20^. The nitrate decline observed in starved culture (Mos) can be ascribed to the heterotrophic denitrification process using organics from cell lysis ^47,48^. Rapid nitrate removal in the first 3 days was explained by the existence of reduced sulfur compounds on pyrite surface ^49^, which was in agreement with previous study using *Thiobacillus thioparus* (a bacterium that can only oxidize sulfur compounds) that nitrate removal was only observed in the first few days ^47^, this was also supported by the fact that acid-pretreated pyrite particles had lower reaction rate (0.004 L/(g·d) ^20^, 0.0004 L/(g·d) ^50^). Nitrite accumulation was usually observed in anaerobic pyrite oxidation process ^17,20,50^ and was considered as the major intermediate product ^17,48^, which was attributed to the inhabitation of nitrate on nitrite reductase ^51^ and the fact that nitrate was more preferred electron accepter than nitrite ^52^. Nitrite accumulation occurred in both cultures that with/without EDTA addition (Figure 2b), while the earlier nitrite accumulation peak occurred along with the faster the denitrification rate. Apart from faster nitrite accumulation, higher nitrite concentration was also perceived in Rec than that in Ctrl, which was ascribed to the more nitrate been reduced in Rec. It was noteworthy that even though nitrite accumulation occurred much earlier and intenser in Rec, of which the nitrite concentration also decreased much faster, which was resulted from the nitrate concentration decline and enough electron supply by help of EDTA.

### 2. Biogeochemical model of anaerobic microbial pyrite oxidation

Combining experimental results and previous studies, a conceptional biogeochemical model of anaerobic microbial pyrite oxidation was proposed (Figure 6), which may provide insights into the couplings of geochemistry, mineralogy, and microbial dynamics within this process. Three main stages were included in the conceptional model, the first stage came the period that microbes colonized at pyrite surface and tend to form localized colonies, during which pyrite was oxidized mainly by direct microbial attack; once simple biofilm formed on pyrite surface, the microbes produced ligands can help pyrite oxidation by chelating with Fe^3+^; finally came complex microbial communities on pyrite surface, and oxidation occurred mainly by soluble Fe^3+^ within acidic micro niche.

**Figure 6.**
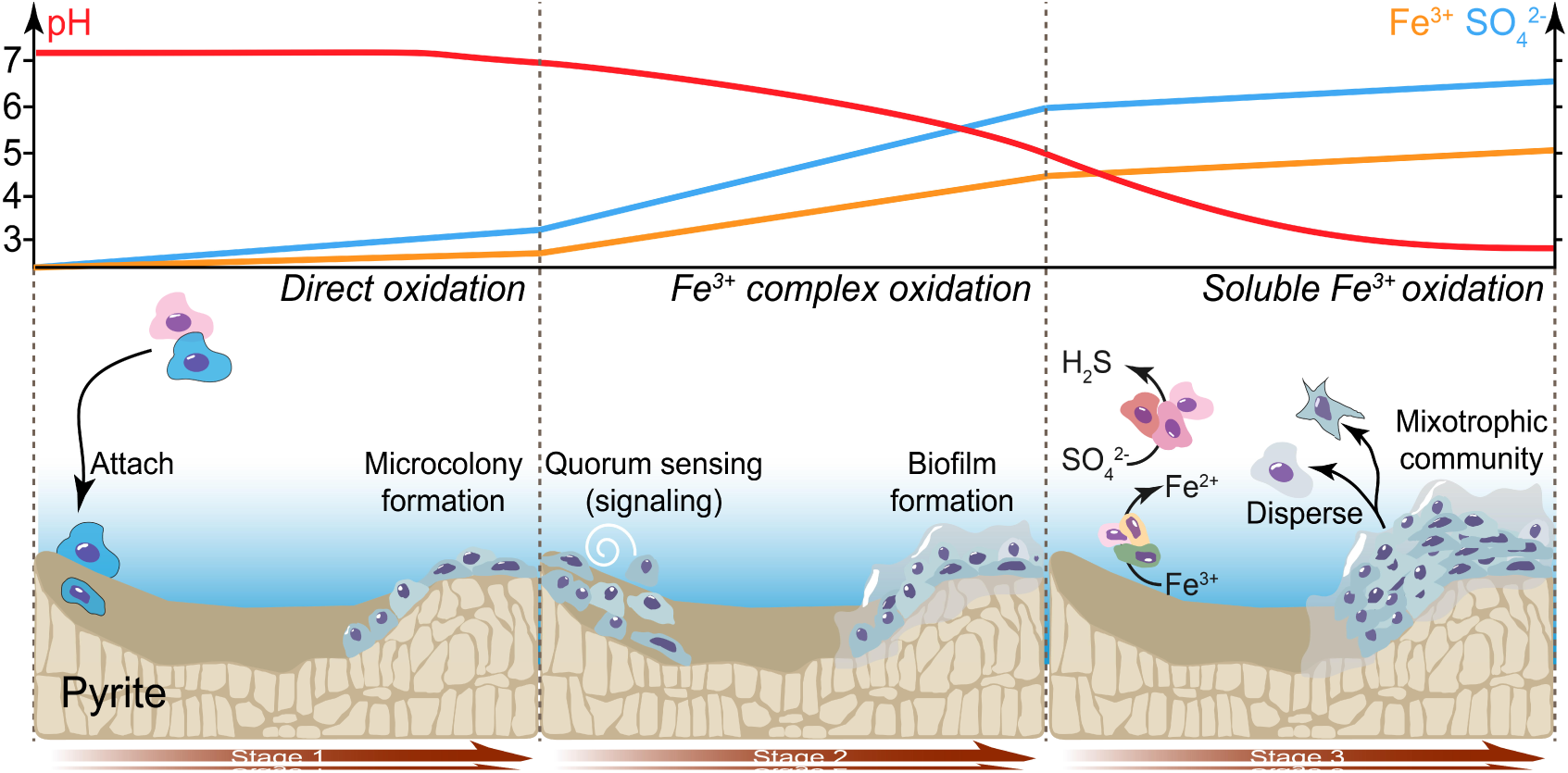
Concept model of anaerobic pyrite oxidation process.

Redox potential is a key factor influencing the initial pyrite oxidation at the first stage. The pyrite oxidation under anaerobic circumneutral conditions can be achieved with potential higher than 200 mV (Figure 1b), which differed from that needed under aerobic conditions (650 mV) ^46^. Thus cellular components with high redox potential could oxidize pyrite by direct attack (Figure 1b). Owing to the deficient energy supply, the energy metabolism was insufficient for ligands production, and inadequate to uphold indirect pyrite oxidation pathway. Considering Fe^3+^ tended to precipitate under neutral pH, pyrite oxidation may proceed in direct pathway (Figure 3), which was supported by the fact that microbes tended to form biofilm on pyrite surface ^53^ and direct cell attachment was always required for pyrite oxidation ^54^. In addition, initial pyrite oxidation happened by surface Fe^2+^ oxidation rather than sulfur oxidation ^28^. Even though sulfur could supply more electrons than Fe^2+^, iron oxidizing microbes may prevail rather than sulfur oxidizing ones. Specifically, for *T*.*denitrificans* (an iron and sulfur oxidizing bacterium) and *Sulfurimonas denitrificans* (*S*.*denitrificans*, a sulfur oxidizing bacterium), a higher abundance for *T*.*denitrificans* in EDTA-addition culture and for *S*.*denitrificans* in EDTA-free culture were therefore reasonable (Figure 4b), which was in consistent with previous column study using different ratio of pyrite and sulfur ^55^. While *Acidovorax sp*. BoFeN1 (an iron oxidizing bacterium that cannot oxidize sulfur) was proved cannot mediating pyrite oxidation ^49^, possible explanation was that, without using sulfur released from pyrite oxidation, electrons from Fe^2+^ oxidation were too few to support cell proliferation.

Along with direct pyrite oxidation by high redox potential cellular compounds, reduced sulfur components that released from pyrite crystal were utilized by microbes and biofilm evolved on pyrite surface ^53^, and then simple microbial communities developed^56^, which was a milestone indicating the second reaction stage. In cultures adding EDTA, high diversity of energy metabolism was observed (Figure 5b), and was confirmed by determination of electron transport system activity (ETSA, Figure S6), reactive oxygen species (ROS, Figure S5), and cytochrome C (Figure S7), which can be attributed to the energy supply increment by utilizing the released reduced sulfur. Microbes with vigorous metabolism had high abundance of transport protein (Figure 5e, g), which was also confirmed in 3D-EEM analysis (Figure 3a). Meanwhile, limited pyrite surface area hindered the ever-increasing cell-mineral attachment (Figure 3c, d), thus microbes may use Fe^3+^ complex with produced ligands as pyrite oxidants preferably in the local reaction region formed by biofilm ^57^. The real process can be similar with cultures adding trace EDTA, within which observed significant pyrite oxidation enhancement (Figure S2). Furthermore, considering the rapid increase of pyrite E_ocp_ with pH drop (Figure 1b), it was not likely for microbes to maintain an acidic microenvironment during this period. It should be noted, owing to the readily available reduced sulfur on pyrite surface ^49^, experiments using crushed pyrite particles may switched into stage 2 shortly, leading to a fast reaction rate initially (Figure 2); on the contrary, pretreatment removing surface oxide layer may result in a reduced reaction rate and sulfate concentration ^20,50,58^, which was an indication for stage 1.

As a result of continuous pyrite oxidation, physicochemical properties of the cell-pyrite surface changed. Key parameters including nitrate, sulfate, and pH may show decrease, increase, and drop tendency, respectively, which can be ascribed to the nitrate consumption and sulfate production mediated by cellular metabolism. Corresponding to physicochemical variation, the surface microbiomes changed with two major attributes, i.e. the strengthened biomass production powered by enough energy supply, and microbial functional differentiation drove by discrepant environmental factors. For cultures adding EDTA, high level phylogenetic diversity (Figure 4b) and functional diversity (Figure 5) was observed on account of sufficient electron supply. In analogue to this stage, in the later period of aerobic pyrite oxidation, the abundance of sulfate reducing bacteria (SRB) that use organics from autotrophic bacterium’s permeation or lysis as carbon source increased ^59^, indicating an ascended community diversity. Since efficient pyrite oxidation can be hardly completed with Fe^3+^ complex (Figure 1b, e), using soluble Fe^3+^ as oxidant may be preferable for microbes under acidic environment, namely the micro-environment would switch from neutral to acidic.

### 4. Environmental implications

Microbial mediated anaerobic pyrite oxidation has important implications for geochemistry on modern earth. For pyrite bearing sediments and shallow aquifers, the penetration of nitrate from various sources may initiate anaerobic pyrite weathering ^17^. Traditionally, a direct chemical reaction between nitrate and pyrite was not expected to occur or examined as very slow kinetics ^60^. Even though the formed nitrite from nitrate radiolysis was capable oxidizing pyrite chemically ^19^, its impact was limited because of the slow reaction kinetics ^61^. However, taking fact that the electrochemical oxidation threshold of pyrite at circumneutral conditions (ca. 200 mV, Figure 1b) was much lower than the pre-assumed value (ca. 650 mV, ^46^), the potential rates and impacts of pyrite oxidation at initially neutral-pH systems should be reexamined, and the importance of microbes should be emphasized since either high redox potential cell compounds or produced organic ligands do accelerate pyrite oxidation. With microbial mediated pyrite oxidation caused by nitrate plume leaching, the mobility of redox-sensitive radionuclides (such as Se, Tc, U, Np, and Pu) ^62^, as well as toxic metals ^63^, may increase, and thereby threaten the eco-security.

Microbial mediated anaerobic pyrite oxidation could also play a role in pollution control, including nitrate contamination attenuation in groundwater, advanced nutrients removal in municipal plants, and mobilization of toxic metals ^20,23,50^. Even though prior studies has focused on nutrients removal in artificial wetlands with pyrite ^24^, pH drop-down in pyrite-based denitrification column ^26^, and immobilization of toxic metals using particular pyrite ^64^, a lack of fundamental understanding of how anaerobic pyrite oxidation occurred has hindered further optimization for most systems. With the provided electrochemical oxidation threshold (Figure 1) and concept model considering different oxidation stages (Figure 6), further guidance for anaerobic pyrite oxidation-based pollutant removal processes was expected.

Furthermore, subglacial microbial community driven by anaerobic pyrite oxidation has been confirmed by meta-transcriptomics ^65^ and sulfur isotope data ^66^, and the existence of active subglacial sulfur metabolism and its possible acceleration for pyrite weathering has been suggested in pyrite bearing sites ^67^. Thus, the concept model proposed here may provide helpful insights into how subglacial microbial community sustained and evolved with a little energy available during glacial-interglacial cycles.

## Materials and methods

Electrochemical analysis of pyrite oxidation under anaerobic circumneutral conditions were conducted with standard three-electrode configuration and pyrite cubic working electrodes cut from natural pyrite minerals. Microbial mediated anaerobic pyrite oxidation experiments were carried out with three main microbial treatments considering microbes under starved (Mos), pyrite (1.0 g/L) addition (Ctrl), and both pyrite (1.0 g/L) and EDTA (5.0 mM) addition (Rec) conditions. Detailed methods are provided in the *SI Appendix*.

## Acknowledgement

The authors acknowledge financial support from the National Natural Science Foundation of China (NSFC) (No. 51578519), the Major Science and Technology Program for Water Pollution Control and Treatment (No. 2017ZX07202002), and the Fundamental Research Funds for the Central Universities (No. 2652018204).

## Supporting Information

The supporting information includes text, figure S1-S12, table S1-S3 and dataset S1 for the detailed materials and methods, detailed experimental design, reproduction of the main experiment, and analysis of related additional experiments.

